# Surviving the Storm: Exploring the Role of Natural Transformation in Nutrition and DNA Repair of Stressed *Deinococcus radiodurans*

**DOI:** 10.1101/2024.07.11.603131

**Authors:** Dhirendra Kumar Sharma, Ishu Soni, Gagan D. Gupta, Yogendra Singh Rajpurohit

## Abstract

*Deinococcus radiodurans*, a natural transformation (NT) enabled bacterium renowned for its exceptional radiation resistance, employs unique DNA repair and oxidative stress mitigation mechanisms as a strategic response to DNA damage. This study excavate into the intricate roles of NT machinery in the stressed *D. radiodurans*, focusing on the genes *comEA*, *comEC*, *endA*, *pilT* and *dprA*, which are instrumental in the uptake and processing of extracellular DNA (eDNA). Our data reveals that NT not only supports the nutritional needs of *D. radiodurans* under stress but also have roles in DNA repair. The study findings establish that NT-specific proteins (ComEA, ComEC, and EndA) might contribute to support the nutritional requirements in unstressed and heavily DNA-damaged cells while DprA contribute differently and in a context-dependent manner to navigating through the DNA damage storm. Thus, this dual functionality of NT-specific genes is proposed to be one of factor in *D. radiodurans* remarkable ability to survive and thrive in environments characterized by high levels of DNA-damaging agents.

**Author Summary:** *Deinococcus radiodurans*, a bacterium known for its extraordinary radiation resistance. This study explores the roles of natural transformation (NT) machinery in the radiation-resistant bacterium *Deinococcus radiodurans*, focusing on the genes *comEA*, *comEC*, *endA*, *pilT*, and *dprA*. These genes are crucial for the uptake and processing of extracellular DNA (eDNA) and contribute to the bacterium nutritional needs and DNA repair under stress. The findings suggest that the NT-specific proteins ComEA, ComEC, and EndA may help meet the nutritional needs of unstressed and heavily DNA-damaged cells, whereas DprA plays a distinct role that varies depending on the context in aiding cells to cope with DNA damage. The functionality of NT genes is proposed to enhance *D. radiodurans* survival in environments with high levels of DNA-damaging agents.

## Introduction

Natural transformation (NT) in bacteria, a form of horizontal gene transfer, is a genetically regulated process that unfolds in an orderly fashion [1]. Its primary aim is to acquire extracellular DNA (eDNA) from environment for various purposes including genome evolution [2–4], DNA repair [5–7], and fulfilling nutritional needs [8, 9]. NT is a driving force in bacterial evolution [10–12], yet its effects on genome stability and removal of deleterious genetic elements remain contentious [13–16]. Research into NT across different bacterial species highlights its varied effects on adaptive evolution, suggesting a multifaceted role that can be advantageous [16, 17], indifferent [18], or contingent on circumstances [17, 19–21]. The direct beneficial impacts of DNA uptake on individual cells could be nutrient source such as nitrogen, carbon, and nucleotides [10, 12, 21] or a template for DNA repair mechanisms [5–7], with experimental findings corroborating these hypotheses across bacterial diversity. The acquisition of genetic material from external environment via NT may carry risks, including the burdens of replication, transcription, and metabolism posed by new genes, as well as potential disruptions to regulatory and protein interaction networks [22]. Additionally, recipient cell faces significant concern regarding the acquisition of genomic parasites, which all self-replicating cell genomes must protect against [23].

*D. radiodurans*, a radioresistant bacterium and this remarkable radioresistance arises from a synergy of multiple strategies. These include efficient repair of DNA double-strand breaks [24–37], protection of proteins from oxidation [38–42], antioxidants protection [31, 38, 43–63], novel DNA repair proteins [64–75], cellular signaling constitute serine/ threonine kinase and cell division regulation [47, 76–89], and a compact nucleoid structure [64, 88, 90, 91], all working together to ensure cell survival after exposure to extremely high doses of gamma rays, prolonged desiccation, and other DNA damaging agents. Additionally, *D. radiodurans* maintains its competency throughout the exponential growth phase, experiencing a decline in transforming frequency only during the stationary phase [92]. DNA transport machinery is conserved in *D. radiodurans*, and a model for the NT of *D. radiodurans* has been proposed [93]. This model describes the initial binding of eDNA with type IV pilus (Tfp) proteins. The DNA uptake machinery is similar to that in *Vibrio cholerae*, comprising homologues of pilins PilIV (DR0548 and DR1232), outer membrane channel PilQ (DR0774), ATPases PilB (DR1964) and PilT (DR1963), and pre-pilin peptidase PilD (DR2065). Additionally, it includes a hypothesized inner membrane channel analogous to PilC in *V. cholerae* [94]. External DNA traverses the outer membrane via the PilQ channel, facilitated by the DNA-binding protein ComEA (DR1855) or ComEA like protein (DR0207), which draws the DNA into the periplasm. Subsequently, periplasmic Endonuclease A (EndA; DR1600) convert the dsDNA to ssDNA by degrading one DNA strand and only one strand of DNA is translocated into the cytoplasm through the ComEC (DR1854) (inner membrane channel), likely assisted by ComF (DR1389). Once inside, the transforming single-stranded DNA (ssDNA) is protected by DNA binding proteins such as single stranded DNA binding (SSB) protein, DdrB, and DprA (DR0120). Differential processing occurs between plasmid and genomic DNA: DprA, bound to genomic DNA, promotes RecA loading onto the genomic DNA, initiating homologous recombination with the host cells genome, while DdrB or RecO, independently of RecA, facilitates enabling of plasmid DNA circularization and establishment through its single-strand annealing (SSA) activity [30, 93, 95]. RecO protein serves as a versatile transformation factor, potentially replacing DdrB for plasmid DNA fragment annealing or substituting SSB/DprA for RecA loading onto genomic DNA, aiding in recombination through the RecFOR pathway [93, 96]. The experimental validation confirmed the significance of various proteins in NT, revealing that gene mutants of Tfp proteins (*ΔpilQ, ΔpilIV, ΔpilD, ΔpilT,* and *ΔpilB*), along with genes encoding proteins responsible for DNA transfer through the cytoplasmic membrane (*ΔcomEC* and *ΔcomEA ΔcomEC*), exhibited a complete absence of transformation, except for *ΔcomF* which demonstrated approximately a 2,000-fold reduction in transformation efficiency [93]. Moreover, proteins like DprA, DdrB, and RecO, crucial for the terminal stage of NT, displayed redundant roles in the transformation of *D. radiodurans*, as only the double mutants *ΔdprAΔddrB* and *ΔdprAΔrecO* exhibited a total loss of transformation [93, 97].

Together, report suggested that Tfp, ComEC-ComEA and DprA plays a crucial role in natural transformation by binding of eDNA to transfer through cytoplasmic membrane, safeguarding, and assisting the integration of incoming DNA into the bacterial chromosome respectively [93]. Additionally, we have recently reported on the roles of DprA in ssDNA binding, which prevents degradation by nucleases [63]. Furthermore, DprA has been shown to play crucial roles in the differential coordination of DNA double-strand breaks (DSBs) repair pathways, contributing to gamma radiation survival in *D. radiodurans* [24]. To explore deeper into the specific roles of NT genes (*pilT, endA, comEA, comEC,* and *dprA*) in surviving the onslaught of DNA damage, we assessed the survival and capacity for repairing DSBs in these genes mutant background. Our findings establish that NT-specific genes might contribute differently and in a context-dependent manner to navigating through the DNA damage storm. Additionally, our findings indicate that eDNA playing a role in providing nutritional support to stressed cells, yet utilizing eDNA for DNA repair appears to detrimental or inconclusive consequences.

## Results

### 1. Natural transformation specific genes mutant showed differential cell survival response to gamma radiation and Mitomycin C

NT integrates eDNA into bacterial genomes through a multi-step process where various NT-specific genes contribute toward internalization of incoming eDNA. This transfer is facilitated by specific proteins such as ComEA, ComEC, ComF, and EndA [98–100]. Importantly, translocated DNA is being protected from cytoplasmic nuclease degradation by proteins like DprA and SSB [101]. To understand the roles of NT-specific genes in cell survival of stressed *D. radiodurans* mutants of NT-genes (*ΔpilT, ΔendA, ΔcomEA,* and *ΔdprA*) were generated and their cell survival monitored against gamma radiation and Mitomycin C (MMC). The *ΔcomEA* and *ΔpilT* gene mutants exhibited cell survival similar to wild type when exposed to various doses of gamma radiation (Fig. 1A). On the other hand, the *ΔendA* mutant showed decrease in the cell survival by one log-cycle while *ΔdprA* showed decrease in cell survival at higher doses of gamma radiation (>12kGy) (Fig. 1A). The MMC had a profound effect on the *ΔendA* mutant (∼1.6 log cycle drop) while *ΔcomEA*, *ΔpilT*, and *ΔdprA* mutants survival nearly equal to wild-type *D. radiodurans* when treated with MMC (10 µg/ml) for 30 min (Fig. 1B). The trans-complementation of *endA* and *dprA* gene from pRAD plasmid (a constitutive expression) in *ΔendA* and *ΔdprA* mutants restore the wild-type phenotype respectively (data not shown). This results suggestive of observed phenotype of *ΔendA* and *ΔdprA* mutants in response to gamma radiation and MMC is effect of absence of respective gene per se instead any polar effect.

**Figure 1:**
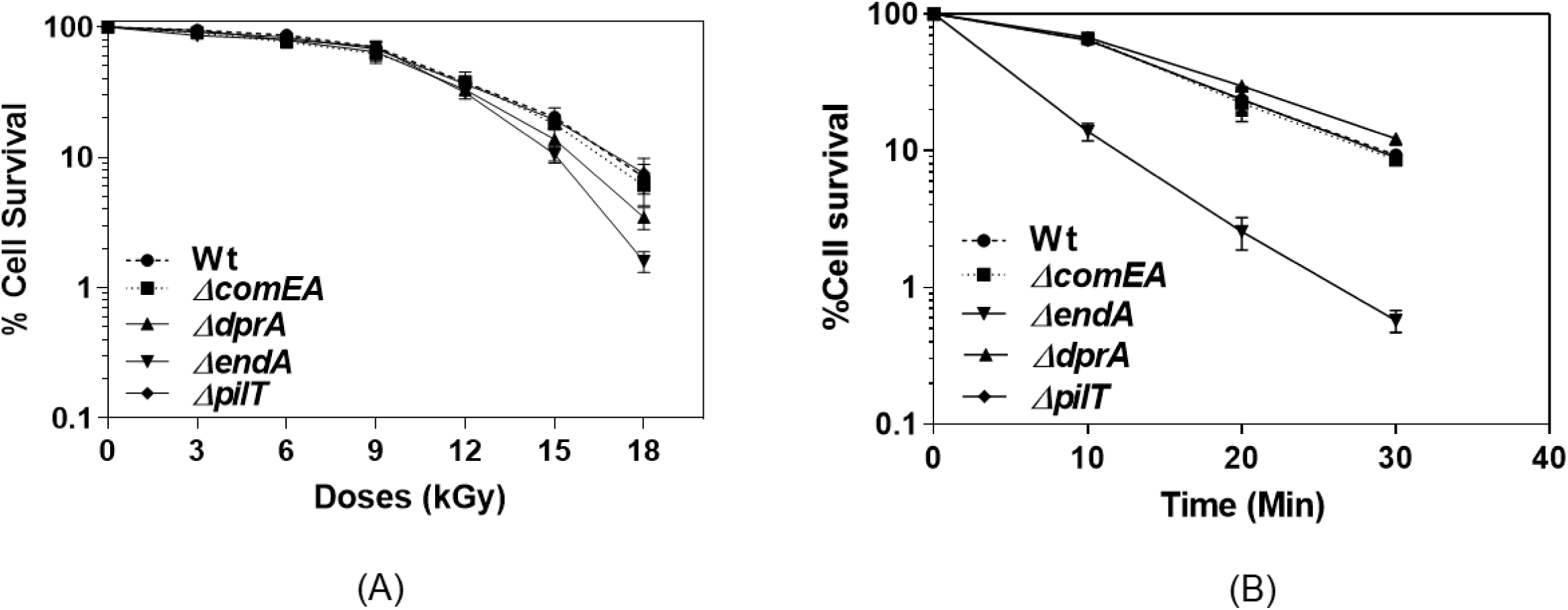
The cell survival of wild type and natural transformation (NT) mutants of *D. radiodurans* after exposure to gamma radiation and MMC. (A) Exponential growth phase cells of wild type (-●-), *ΔcomEA* (-▪-), *ΔendA* (-▾-), *ΔdprA* (-▴-), and *ΔpilT* (-♦-) mutants were subjected to different doses of gamma radiation, and the percentage of cell survival fraction was plotted as a function of the gamma radiation doses (kGy). The mean ± SEM of two independent experiments were shown. (B) Exponential growth phase cells of wild type (-●-), *ΔcomEA* (-▪-), *ΔendA* (-▾-), *ΔdprA* (-▴-), and *ΔpilT* (-♦-) mutants were exposed to MMC (10µg/ml) doses ranging from 0 to 30 minutes. After dilution, plated on TGY medium and incubated at 32^0^C for 48 hours. The percentage of cell survival fraction was plotted as a function of the MMC treatment time.The mean ± SEM of two independent experiments were shown.

Furthermore, the growth pattern of unstressed wild-type and different mutants (*ΔendA*, *ΔcomEA*, and *ΔdprA*) was nearly identical to wild type albeit the *ΔendA* mutant showed slight slow growth pattern, indicating that genes (*ΔcomEA*, and *ΔdprA*) deletions had no effect on the normal growth while *ΔendA* gene deletion slowdown the growth slightly of *D. radiodurans* (Fig. 2A). However, when cells were exposed to 6kGy doses of gamma radiation *ΔendA* mutant exhibited some growth defects (Fig. 2B). Furthermore, growth defect effect of *ΔendA* mutant become more profound at gamma doses of 12kGy and showed ∼600 minutes long lag phase compare to wild type (∼400 minutes) (Fig. 2C). Interestingly, at 12kGY dose *ΔdprA* mutant also showed growth defect (Fig. 2C). Together, the observation of long lag phase post gamma irradiation indicate the defect in DNA repair potential of *ΔendA* and *ΔdprA* mutants. The growth pattern of *ΔcomEA* under gamma radiation stress is similar to that of the wild-type (Fig. 2B & C). This suggests that ComEA may not play a significant role in radiation resistance, or that the *comEA* homolog, *dr0207*, could be compensating for the loss of ComEA function.

**Figure 2:**
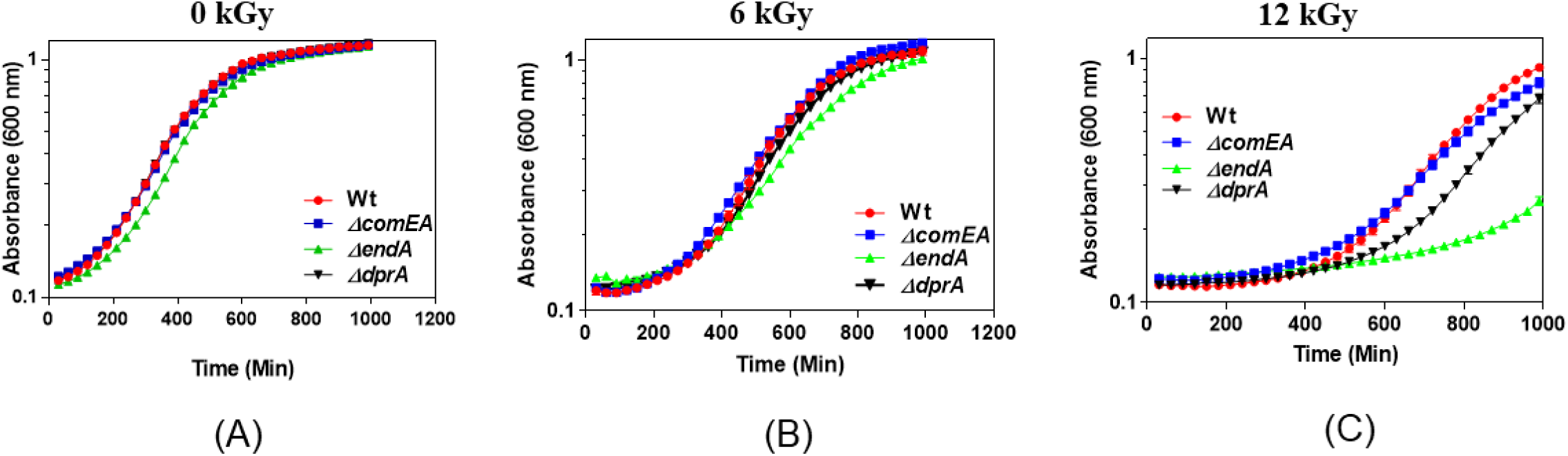
Cell viability and growth curve of wild type and its NT mutants after exposure to gamma radiation. Optical density at 600 nm was measured continuously using a microtitre-based density reader, and the growth medium (TGY) served as a blank for the entire incubation period. The obtained data was normalized with the blank optical density. Panel (A) shows the growth of normal, untreated cells, panel (B) displays cells treated with 6 kGy of gamma radiation, and panel (C) illustrates cells treated with a dose of 12 kGy of gamma radiation.

### 2. Analysis of DSB repair profiles in *D. radiodurans* NT genes mutant using pulse field gel electrophoresis (PFGE)

In order to investigate the DSB repair profiles of *ΔcomEA*, *ΔdprA*, and *ΔendA* mutants, pulse field gel electrophoresis (PFGE) was employed. The *ΔdprA* mutant showed faster DSB kinetics to repair DSBs introduced by gamma radiation (6kGy) (Fig. 3A), as reported recently [24]. The *ΔcomEA* mutant exhibited a DSB repair pattern identical to that of the wild-type (Fig. 3B). On the other hand, the *ΔendA* mutant showed slightly slower DSB repair compared to the wild type when exposed to 6kGy gamma radiation (Fig. 3B). The slow DSB repair is evident at 4 hours post irradiation time (PIR) in the *ΔendA* mutant, as a higher molecular weight repaired genome in the *ΔendA* mutant is missing at 4 hours PIR time (Fig. 3B). However, the wild type and *ΔcomEA* mutant showed nearly fully repaired genomes at 4 hours PIR (Fig. 3). Results suggest that *ΔcomEA* function is not necessary or in its absence its functional homolog (*dr0207*) might compensate *ComEA* functions for DSB repair, while *ΔendA* might play a role in the repair of DSBs induced by gamma radiation. Interestingly, the *ΔdprA* mutant showed faster DSB repair compared to the wild-type, and within 1 hour of post-irradiation recovery (PIR) time, distinct repaired bands appeared on the gel (Fig. 3A). Together, these findings indicate that the *dprA* and *endA* genes are involved in DSB repair, and their absence results in faster or slower repair kinetics, respectively. It is worth noting that both the *ΔdprA* and *ΔendA* mutants were sensitive to gamma radiation at higher doses, although the level of gamma sensitivity varied (Fig. 1A). Despite this, the DSB repair kinetics were different, with the *ΔendA* mutant exhibiting delayed DSB repair (Fig. 3B) and the *ΔdprA* mutant showing faster DSB kinetics (Fig. 3A). This suggests that the mechanisms of DSB repair induced by gamma radiation may differ in these mutants. Furthermore, the PFGE banding pattern for all NT-specific genes was nearly identical to that of the wild-type, indicating that there were no large genomic rearrangements (Fig. 3). Unlike with gamma radiation, the *ΔendA* mutant displayed significant sensitivity when treated with MMC (Fig. 1B). This sensitivity in the *ΔendA* mutant is due to defective DSB repair, as no detectable DSB repair was observed 6 hours post-MMC treatment (Fig. 4B). However, other NT-specific mutants, such as *ΔdprA* and *ΔcomEA*, exhibited DSB repair profiles similar to the wild type, although the *ΔcomEA ΔcomEC* mutants showed relatively slower DSB repair (Fig. 4).

**Figure 3:**
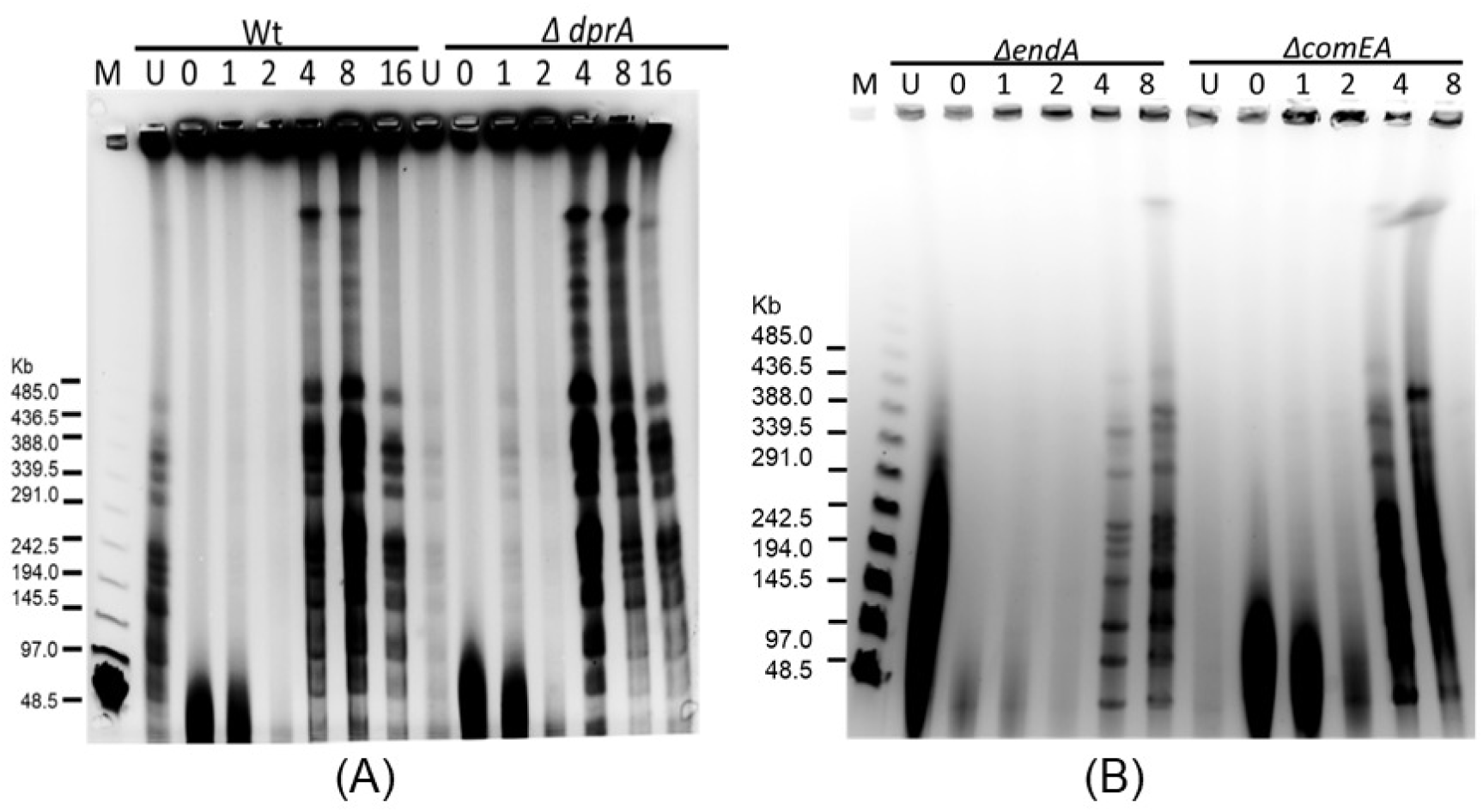
The kinetics of repair of DNA double-strand breaks (DSBs) in wild-type and NT mutants of *D. radiodurans*. Pulsed-field gel electrophoresis (PFGE) was utilized to evaluate the repair kinetics. The NotI-digested DNA from unirradiated cells (U) and irradiated cells at different post-irradiation time points (PIR) after exposure to 6kGy were visualized immediately after irradiation (0) and at specified incubation times (hours) on the gel. Lambda PFG molecular mass standards (lane-M). (A) The kinetics of DSB repair in wild-type (Wt) and *ΔdprA* mutants are shown, while (B) displays the kinetics of DSB repair in *ΔendA* and *ΔcomEA* mutant.

**Figure 4:**
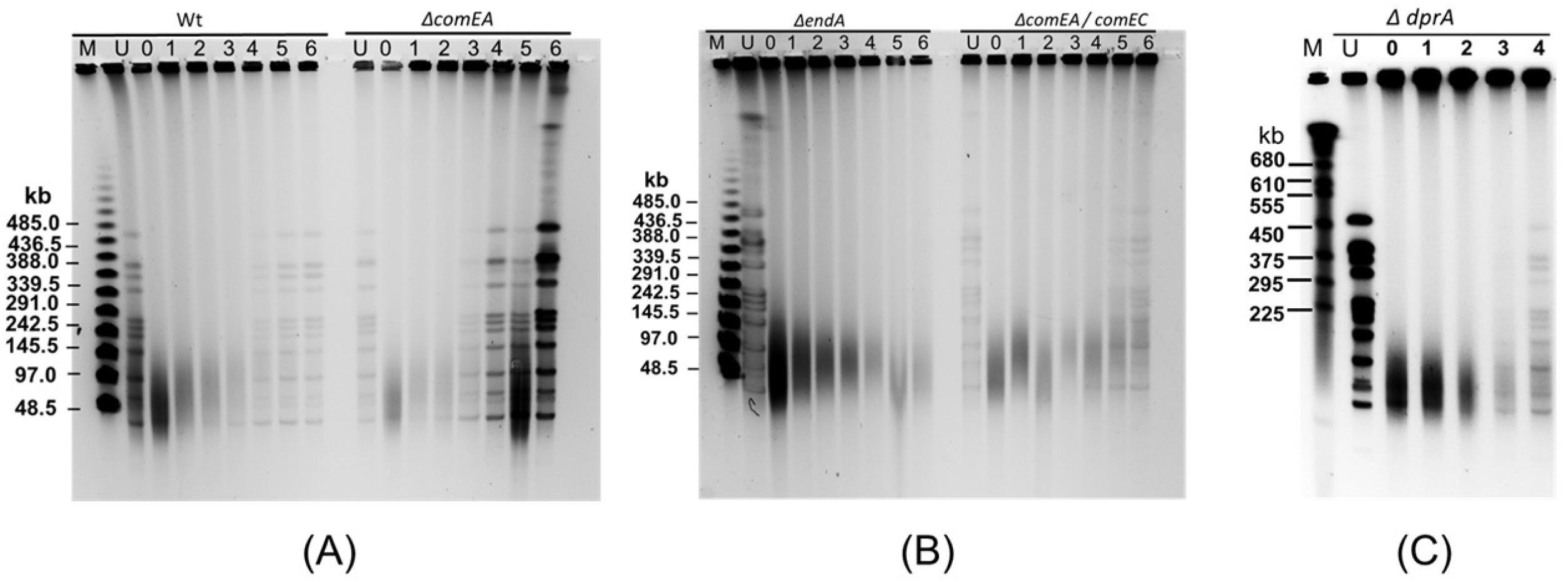
DNA Double-Strand Break (DSB) repair kinetics in *D. radiodurans* wild-Type and NT mutant strains treated with MMC. The figure illustrates the kinetics of DSB repair in wild-type and four NT-specific mutant strains of *D. radiodurans*—namely, wild type (Wt), *ΔcomEA*, *ΔendA*, *ΔcomEAΔcomEC*, and *ΔdprA*—following treatment with mitomycin C (MMC). The repair kinetics were assessed using pulsed-field gel electrophoresis (PFGE). NotI-digested DNA samples from untreated cells (U) and cells treated with 10 µg/ml MMC for 30 minutes were collected at various recovery time points (4 to 6 hours post-treatment, PTR) in TGY medium. These samples were analyzed by PFGE to evaluate the DSB repair process. Lane-M contains molecular mass standards for reference.

In summary, the findings suggest that in NT-specific gene mutants, DSBs induced by gamma radiation may follow wild-type similar DSB repair kinetics (*ΔcomEA*), fast repair kinetics (*ΔdprA*), or slow/equal rate repair kinetics (*ΔendA*). In contrast, MMC-induced DSBs follow comparable repair kinetics of wild type and all tested NT mutants except *ΔendA* mutant. These observations indicate the possibility of differential utilization of DNA damage repair machinery to repair gamma radiation and MMC-induced DNA damage in NT gene mutants. Furthermore, these observations highlight the complexity of DSB repair mechanisms in *D. radiodurans*.

## 3. ΔendA and ΔdprA single or ΔendA ΔdprA double mutants exhibited differential responses to DNA damage induced by combined gamma radiation+UV or MMC+UV treatments

Data from preceding section suggested that the *ΔcomEA* and *ΔpilT* mutant strains did not show significant differences in cell survival compared to the wild-type strain under all tested conditions, indicating that these genes may not be necessary for *D. radiodurans* to cope with DNA damage (Fig. 1). The *ΔendA* mutant strain showed decrease in survival to gamma radiation and MMC treatment, suggesting that the *endA* gene plays a role in coping with gamma radiation and MMC induced DNA damage (Fig. 1, 3, 4 & S1). In contrast, the *ΔdprA* mutant strain showed a susceptibility to gamma radiation at doses beyond 10kGy and showed nearly similar survival after MMC treatment (Fig. 1 & S1), indicating that the *dprA* gene is crucial for coping with high doses of gamma radiation. Studies found that both gamma radiation and MMC primarily cause DSBs to the genome, but the mechanisms of DSB generation differ between the two agents and additionally gamma radiation introduce oxidative stress and ssDNA breaks while MMC also damage proteins and RNA [102–104]. The gamma radiation causes the DSBs by direct deposition of energy to phosphodiester bond or indirectly through reactive oxygen species and other radicals. MMC causes the inter and intra-strand cross links in DNA and these cross-links seldom converted to DSBs through DNA replication [105, 106]. Thus, it could be conceivable that mechanism of biomolecule damage and DSBs generation is different for the gamma radiation and MMC [57, 103, 107, 108].

The contrasting responses of *ΔendA* and *ΔdprA* mutants to MMC and gamma radiation are intriguing, given that both treatments induce DSBs and oxidative damage. The NT-specific EndA nuclease has been shown to support nutritional requirements under conditions of nutrient deprivation [109–111] or the dispersion of bacteria from biofilm [112–114]. The decreased cell survival of *ΔendA* mutants against gamma radiation and MMC suggests that the EndA nuclease may support stressed cells in an unknown manner. We hypothesize that the *endA* gene might play a role in providing nutritional support to stressed *D. radiodurans* cells following exposure to gamma radiation or MMC. However, for eDNA to serve as a source of nutrition in the form of nucleosides, it must remain undamaged. To bolster this hypothesis, we treated wild-type and various mutants (*ΔendA*, *ΔdprA*, and *ΔendA ΔdprA*) with gamma radiation or MMC combined with UV radiation to induce complex DSB damage along with UV-induced DNA damage, including cyclobutane pyrimidine dimers (CPDs) and 6-4 photoproducts. This combined treatment could severely damage cellular DNA, including eDNA, compromising its availability for nutrition. Our data indicate that gamma radiation alone (6 kGy) reduced the cell survival of the *ΔendA* mutant in a marginal reduction 0.1 log-cycle, while UV radiation (1500 J/m²) resulted reduction of 0.3 log-cycle (Fig. 5A & S1). The combined effect of gamma radiation and UV drastically reduced the cell survival of both the wild-type and *ΔendA* mutant, whereas for the *ΔdprA* mutant, the combined effect of gamma radiation and UV led to higher survival by approximately 0.6 log cycles compared to the wild-type and *ΔendA* mutant (Fig. 5A & S1). Conversely, the cell survival of *ΔdprA* mutants remained similar to the wild-type at selected gamma radiation doses but declined marginally with MMC treatment (Fig. 5A & B). Furthermore, when these mutants were tested for MMC in combination with UV, the effect of MMC alone was similar for the wild-type and *ΔdprA* mutants, whereas there was a nearly two log-cycle reduction in survival for *ΔendA* mutants (Fig. 5B). The combined effect of MMC and UV severely impacted the survival of the wild-type and *ΔendA* mutant, but the *ΔdprA* mutant showed approximately 2 log cycles better survival. This improved survival could be attributed to DprA role in the differential regulation of DSB repair pathways in *D. radiodurans*, as recently reported by us [24].

**Figure 5:**
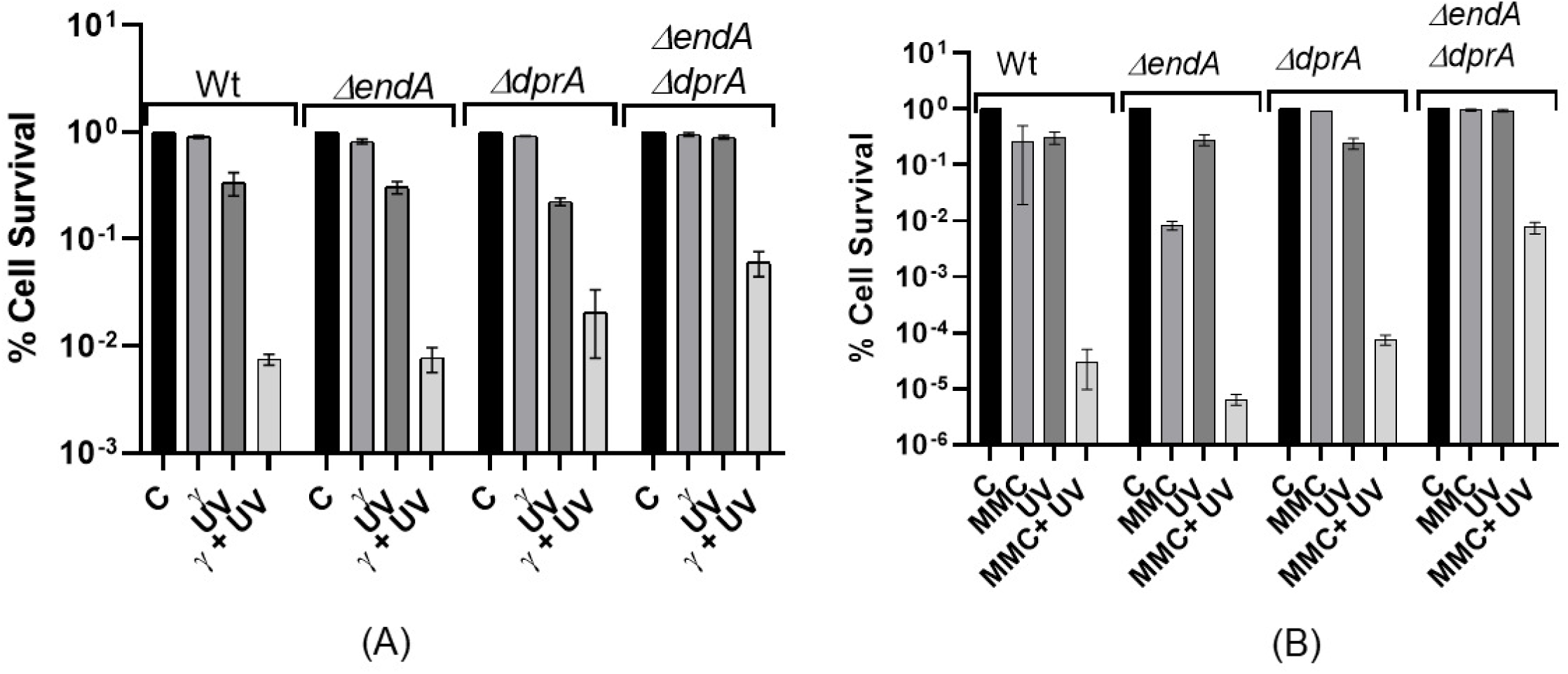
The cell survival of wild-type *D. radiodurans* and various NT mutants were measured after exposure to gamma radiation or MMC combined with UV radiation. The cell survival rates of wild-type *D. radiodurans* and various NT mutants were measured after exposure to gamma radiation or MMC combined with UV radiation. Exponentially growing cells of the *D. radiodurans* R1 wild-type and NT mutants were subjected to (A) 6 kGy doses of gamma radiation, 1500 J/m² UV radiation, or both, and (B) MMC (20 µg/ml) for 30 minutes, 1500 J/m² UV radiation, or both. The percentage of surviving cells was plotted for the different mutants, and the mean ± SEM of two independent experiments was presented.

The *ΔendA* mutant, compromised in NT, and the absence of the *endA* gene led to reduced survival against gamma radiation, MMC, or combined gamma radiation + UV or MMC + UV treatment (Fig. 1, 5 & S1), highlighting the roles of *endA* in coping with genetic perturbations, possibly through its roles in nutritional support during stress (Fig. S2). Furthermore, in the absence of stress, the *ΔendA* mutant growth followed the wild-type pattern, suggesting that *endA* may not have roles in unstressed cells (Fig. S2 A). The *ΔdprA* mutant also showed a defect in NT, and its roles in dealing with gamma radiation and MMC-induced DNA damage are intriguing (Fig. 5). This reflects both its roles in NT and its differential effect on DSB repair pathways such as extended synthesis-dependent strand annealing (ESDSA) and single-strand annealing (SSA) as suggested recently [24].

Interestingly, similar to the *ΔdprA* mutant, *ΔendA ΔdprA* double mutant showed significantly improved cell survival compared to the individual wild-type when treated with gamma radiation + UV or MMC + UV (Fig. 5A, B) and showed better DSB repair kinetics when treated with gamma radiation (Fig. S3). Additionally, this effect is supported by growth curve analysis of *ΔendA ΔdprA* double mutant treated with both gamma radiation and MMC (Fig. S2). The *ΔendA ΔdprA* double mutant showed the shortest lag phase and started to recover faster than the wild-type or individual mutants of *endA* and *dprA* genes (Fig. S2 D), corroborating the findings of Fig. 5. Again, this improved survival could be attributed to DprA role in the differential regulation of DSB repair pathways in *D. radiodurans*, as recently reported by us [24]. Thus, the hypothesis that severely damaged eDNA is less utilized for nutritional purposes is strongly supported by data showing a severe reduction in cell survival of *ΔendA* mutant when gamma radiation or MMC treatment is combined with UV radiation (Fig. 5A, B). Additionally, our findings support previous observations that *D. radiodurans* continuously acquires eDNA during its growth. It can repair UV radiation-damaged transforming DNA through its highly effective repair system, even after chromosomal conversion [92, 115].

### 4. NT-specific genes (*endA*, *comEA* and *comEC*) required for the DNA as a nutritional source

Under specific conditions, bacteria have shown the ability to utilize DNA as elemental sources of phosphorus (P), carbon (C), and nitrogen (N) for their growth [109, 116–119]. This is achieved through two mechanisms: either bacteria secrete DNase into the environment, where these nucleases degrade the free-form eDNA into nucleotides or nucleosides that are then utilized by bacteria [117, 120]. Additionally, bacteria also utilize secretory phosphatases to release phosphate from eDNA for subsequent utilization. However, secreting DNase / phosphatases into the environment is not advantageous because bacteria typically coexist with many other bacterial species. The digested DNA monomers and fragments are easily taken up by competing bacteria and cross feeding reduce the survival benefits. Furthermore, the effective concentration of DNase becomes much lower if it is secreted into the culture medium, making it energetically unfavorable for the bacteria. Thus, bacteria adopt another effective, energy-efficient, and competition-free mechanism is to acquire eDNA into the bacterial periplasm and then utilize it for nutrition or as a structural constituent in biofilms with the help of periplasmic DNase and phosphatases [109–111, 114, 116, 121]. The ComEA protein, which is a receiver of transforming DNA in the periplasm, is localized in the periplasm of *D. radiodurans* with high reliability. Bioinformatics analysis using the DeepTMHMM server (DeepTMHMM-1.0) (https://services.healthtech.dtu.dk/service.php?DeepTMHMM-1.0) predicted that ComEA contains a transmembrane helix at the N-terminus of the protein (residues 7-29), with the remainder of the protein (residues 30-130) located in the periplasmic region. In contrast, *D. radiodurans* EndA is predicted to be cytoplasmic and lacks any signal peptide, as analyzed using the SignalP 6.0 server (SignalP 6.0) (https://dtu.biolib.com/SignalP-6). The *D. radiodurans* EndA catalytic core showed substantial sequence similarity with approximately 25% sequence identity and 30% sequence similarity to *Escherichia coli, V. cholerae,* and *Aeromonas hydrophila* EndA (Fig. S4). However, the N-terminal region (residues 1-90) of *D. radiodurans* EndA shows little sequence identity compared to other EndA proteins (Fig. S4). Nonetheless, *D. radiodurans* has a protein transport system to send proteins across the cytoplasmic membrane via the SecA pathway or the signal recognition particle (SRP) pathway, and it is shown that many designated cytoplasmic proteins, including catalases (KatE1 and KatE2), are transported to the periplasm using SRP pathways [122]. EndA, which is a designated periplasmic protein in many bacteria including *E. coli* [109], *V. cholerae* [94, 123], and *Streptococcus pneumoniae* [124, 125], could have a signal peptide for periplasmic location or could be transported to the periplasm by various transport systems such as Tat, SecA, or SRP pathways [126, 127]. Since, EndA function has been found to be crucial for *D. radiodurans* cells survival exposed to gamma radiation and MMC (Fig. 1 & 5). We assume that under stressed conditions, *D. radiodurans* can utilize eDNA liberated from either lysed cells or from other cells through active secretion for the purpose of nutrition or for the DNA repair purpose. This eDNA can be transported into the periplasmic space of *D. radiodurans*, where the action of EndA liberates free nucleotides as well as ssDNA from incoming dsDNA. Both free nucleosides or nucleosides and ssDNA (once entered inside the cell) can be further utilized as nutrition sources of phosphorus (P), carbon (C), and nitrogen (N).

We have attempted to test this hypothesis by growing the *D. radiodurans* in the minimal medium where calf thymus DNA (cfDNA) being used as sole carbon (C) or nitrogen (N) source (Fig. S5). Unfortunately, we found poor growth support of *D. radiodurans* in minimal medium with and without cfDNA (0.5mg/ml) added (Fig. S5). To further substantiate the findings we have grown wild-type as well different mutants (*ΔendA*, *ΔdprA*, *ΔcomEA*, and *ΔcomEA ΔcomEC*) in TGY medium or TGY medium supplemented with cfDNA DNA (0.5mg/ml). Data suggested that in TGY medium wild-type and all mutants grow near identical and follow the standard growth trajectories (Fig. 6A). However, in the presence of cfDNA *ΔendA*, *ΔcomEA, and ΔcomEA ΔcomEC* mutants growth slowed down compare to *ΔdprA* and wild-type (Fig. 6B). Interestingly, it is observed that the wild type cells takes ∼400 minutes to reach 0.5 optical density (OD) in TGY medium which increased up to ∼750 minutes in TGY+DNA medium to reach similar OD values (compare Fig. 6A and B). The observed doubling in the time required to reach the exponential phase in the presence of cfDNA DNA suggests that utilizing DNA as a nutritional source may take more time. However, this effect was not seen when the cfDNA was pre-digested with DNase and added to the TGY medium (Fig. S6). Instead, *D. radiodurans* cells grew better when provided with pre-digested DNA in the TGY medium (Fig. S6). This finding indicates that small DNA fragments or nucleosides can be utilized by *D. radiodurans* cells. This notion is further supported by growth data obtained for *ΔendA*, *ΔcomEA*, and *ΔcomEA ΔcomEC* mutants, whether in unstressed conditions (Fig. 6B) or gamma radiation-stressed cells supplemented with DNA (Fig. 6D). It was observed that these mutants, which are defective in receiving eDNA in the periplasm (*ΔcomEA*), or defective in both receiving DNA in the periplasm and transporting it through the inner membrane (*ΔcomEAΔcomEC*), or defective in converting dsDNA to ssDNA in the periplasm (*ΔendA*), exhibited delayed growth (Fig. 6B, D). This delay is likely due to their inability to utilize DNA as a nutritional source under both unstressed and stressed conditions. Together, our data suggested that *D. radiodurans* grows better when small DNA or nucleotides or nucleosides available in medium in both stressed and unstressed condition. Furthermore, mutants lacking specific DNA-processing proteins exhibit delayed growth. This suggests that internal processing of DNA in wild-type is more efficient and advantageous for bacterial survival and growth.

**Figure 6:**
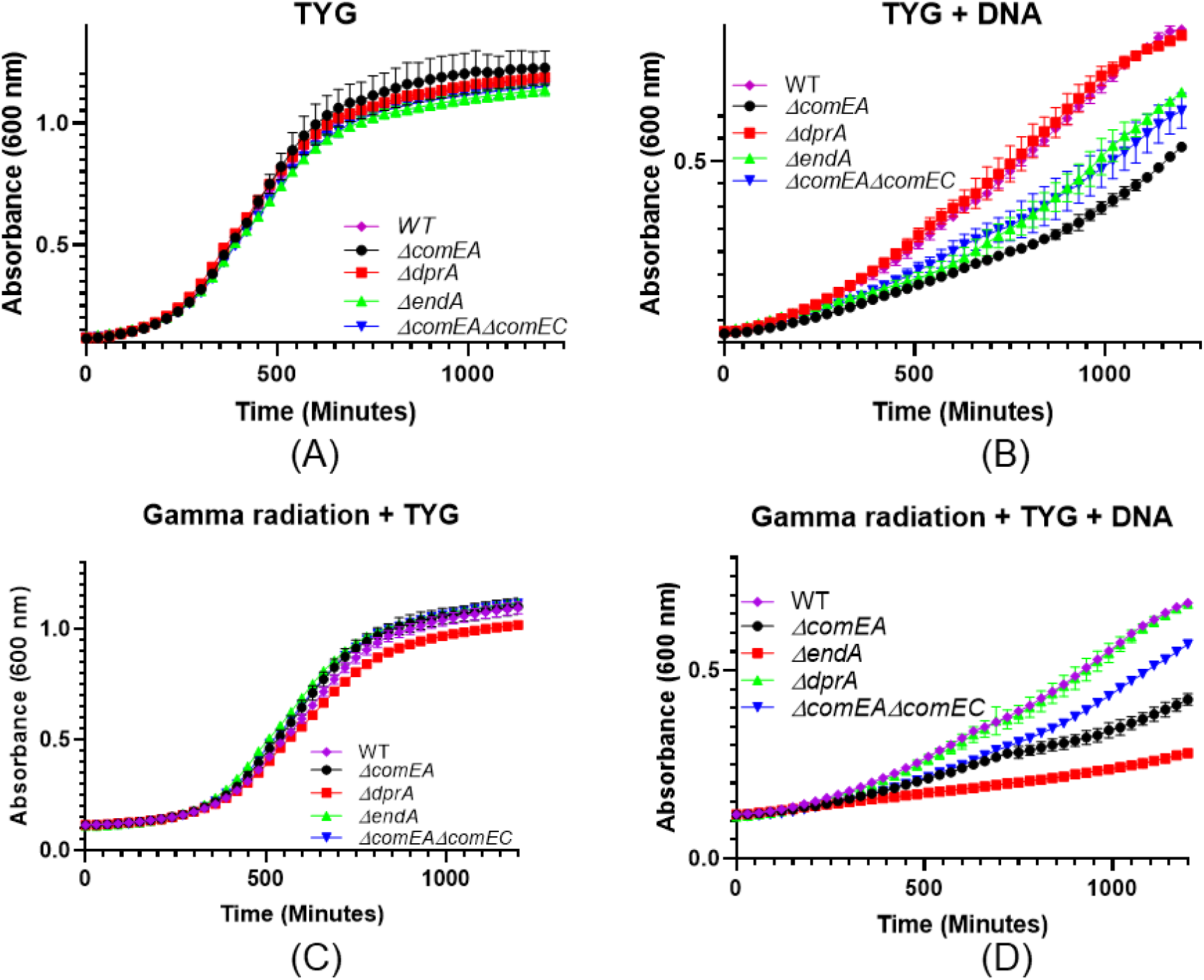
Cell growth curve of wild type and NT mutants after exposure to calf thymus DNA (cfDNA) in TGY growth medium with and without gamma radiation. Optical density at 600 nm was continuously measured using a microtitre-based density reader. Panel (A) shows the growth of normal, untreated cells in TGY growth medium. Panel (B) displays cells treated with cfDNA (0.5 mg/ml) in TGY medium. Panel (C) illustrates cells treated with a dose of 6 kGy of gamma radiation in TGY medium. Panel (D) depicts cells treated with a dose of 6 kGy of gamma radiation in TGY medium supplemented with cfDNA.

## Discussion

The results of this study provide insights into the role of natural transformation (NT) specific genes (*ΔcomEA*, *ΔpilT*, *ΔendA*, and *ΔdprA*) in the survival and DNA repair mechanisms of *D. radiodurans* under various stress conditions, including exposure to gamma radiation, Mitomycin C (MMC), and combinations of gamma radiation or MMC with UV radiation. The first key finding is the differential cell survival response observed among the NT-specific gene mutants when exposed to gamma radiation and MMC. While *ΔcomEA* and *ΔpilT* mutants exhibited survival rates similar to the wild type, the *ΔendA* and *ΔdprA* mutants displayed decreased survival under specific stress conditions (Fig. 1 & S1). Notably, *ΔendA* and *ΔdprA* mutants showed increasing susceptibility to higher doses of gamma radiation (Fig. 1A) while the *ΔendA* mutant showed a significant decrease in survival upon exposure to MMC, indicating a potential role for the *endA* gene in coping with MMC-induced DNA damage (Fig. 1B). Additionally, the growth patterns of the mutants were comparable to the wild type under normal conditions, but defects were observed post-gamma irradiation or MMC treatment, particularly for the *ΔendA* and *ΔdprA* mutants, suggesting impaired DNA repair potential (Fig. 2 & S2).

The analysis of DSBs repair profiles using PFGE further elucidated the roles of NT-specific genes in DNA repair mechanisms. The *ΔdprA* mutant exhibited faster DSB repair kinetics post-gamma irradiation (Fig. 3A), whereas the *ΔendA* mutant showed slower repair compared to the wild type (Fig. 3B). These findings suggest that DprA and EndA are involved in DSB repair, with their absence resulting in altered repair kinetics (Fig. 3). Interestingly, the DSBs repair kinetics of *ΔdprA* mutant differed between gamma radiation-induced DSBs (fast DSBs repair) and those induced by MMC (normal DSBs repair), while *ΔendA* mutant exhibit slow DSBs repair indicating distinct DSBs repair mechanisms for different types of DNA damage and mutant background highlight the complexity of repair followed in *D. radiodurans* (Fig. 3 & 4). Additionally, the observation that the *ΔendA* mutant growth was affected by gamma radiation (Fig. 2C) and MMC (Fig. S2C), resulting in a prolonged lag phase, further supports the notion that EndA plays a role in DNA repair and that its absence leads to a defect in the *D. radiodurans* ability to recover from DNA damage.

Bacteria can utilize DNA as a source of phosphorus, carbon, and nitrogen for growth [109, 116–119]. This is achieved by internalizing eDNA into their periplasm, where periplasmic DNases and phosphatases process it for nutritional or structural purposes [109–111, 114, 116, 121]. The NT-specific EndA nuclease has been demonstrated to aid in nutritional needs during nutrient deprivation [109–111] and in the dispersion of bacteria from biofilms [112–114]. The reduced cell survival observed in *ΔendA* mutants exposed to gamma radiation and MMC indicates that the EndA nuclease might assist stressed *D. radiodurans* cells possibly roles in providing nutritional support to stressed *D. radiodurans* cells after exposure to gamma radiation or MMC. To test this hypothesis, we explored the combined effects of gamma radiation and UV or MMC and UV on cell survival. We considered that combined treatment could severely damage cellular DNA, including eDNA, thereby compromising its availability for nutrition. The *ΔendA* and *ΔdprA* mutants exhibited different responses to these treatments, with the *ΔdprA* mutant showing improved survival when gamma radiation or MMC was combined with UV (Fig. 5). This intriguing finding suggests that DprA may play a role in the differential regulation of DSB repair pathways, especially under conditions of complex DNA damage. These differential roles of DprA could be akin to our recent findings, where we have shown that in the *ΔdprA* mutant, single-strand annealing (SSA) repair assumes a more prominent role in the repair process, along with a modest increase in the function of extended synthesis-dependent strand annealing (ESDSA). Thus, the varying levels of SSA and ESDSA contributions highlight the role of DprA in selecting appropriate double-strand break (DSB) repair pathways in heavily irradiated *D. radiodurans* cells [24]. The *Δend* mutant and the *ΔendA ΔdprA* double mutant, which lacks the ability to undergo natural transformation, also showed distinct responses. The *Δend* mutant demonstrated reduced survival, whereas the *ΔendA ΔdprA* double mutant showed improved survival and faster DSB repair kinetics and growth when treated with gamma radiation + UV or MMC + UV (Fig. 5, S2D, & S3). Collectively, these results support the hypothesis that severely damaged eDNA is less utilized for nutritional purposes, while the possible differential inhibitory effect of DprA on DSB repair pathways leads to elevated cell survival and growth under stress from gamma radiation, MMC, or both agents. We further substantiate the notion that DNA can be utilized as a nutritional resource under stressed conditions, with EndA and ComEA potentially playing roles in this process within the periplasmic space. Our analysis of the growth patterns of the *ΔendA*, *ΔcomEA*, and *ΔcomEA ΔcomEC* mutants in the presence of cfDNA DNA reveals that these mutants are impaired in their ability to utilize DNA as a nutritional source in gamma radiation stressed conditions and unstressed condition, likely due to their inability to process eDNA effectively (Fig. 6).

Together, the findings from this study have significant implications for our understanding of natural transformation roles in DNA repair and cell survival mechanisms in *D. radiodurans*. The differential responses of the *ΔendA* and *ΔdprA* mutants to DNA-damaging agents highlight the complexity of the DNA repair processes in this organism. The observation that the *ΔdprA* mutant exhibits faster DSB repair kinetics is particularly noteworthy. This finding suggests that DprA may be involved in the regulation of DSBs repair pathways of *D. radiodurans*. The role of EndA in the *D. radiodurans* response to DNA damage is also of great interest. The slower DSB repair kinetics observed in the *ΔendA* mutant and the growth defects following gamma radiation / MMC exposure suggest that EndA plays a critical role in the repair of DSBs. The hypothesis that EndA supports stressed cells through nutritional means is strengthen by data provided in this study. However, further research is needed to elucidate the exact mechanisms by which EndA contributes to cell survival under stress by offering nutritional support. The roles of DprA seems to interesting and suggest that relay on eDNA for DNA repair may have detrimental or inconclusive consequences as shown for other bacteria [9, 15, 22, 109, 112, 116]. Thus, the ability of *D. radiodurans* to utilize eDNA for nutrition or DNA repair, whether under unstressed or stressed conditions, likely involves complex dynamics and consequences and repercussions.

## Acknowledgements

The authors would like to express their gratitude to Prof. Fabrice Confalonieri and Prof. Pascale Servant from Université Paris-Saclay, CEA, CNRS, Institute for Integrative Biology of the Cell, France, for kindly providing *D. radiodurans ΔcomEA ΔcomEC* mutants strain.

## Experimental Procedure

### Bacterial strains and plasmids, chemicals, and growth medium

In this investigation, a range of bacterial strains were utilized, including the wild type *D. radiodurans* R1 (ATCC 13939) sourced from the ATCC. The construction of *ΔcomEA, ΔdprA, ΔendA, ΔdprA ΔendA,* and *ΔpilT* mutants was performed in our laboratory using previously described methods [128]. Additionally, *ΔcomEA ΔcomEC* double mutants were obtained from Prof. Pascale Servant, Université Paris-Sud Orsay, France. All strains were cultured in TGY medium (1% Bacto-tryptone, 0.1% glucose, 0.5% yeast extract) with the addition of suitable antibiotics, following the established protocol [86]. The media were supplemented with various antibiotics, specifically kanamycin (8 µg/ml), chloramphenicol (3 µg/ml), and tetracycline (2 µg/ml) for *D. radiodurans*, and ampicillin (100 µg/ml) for *E. coli*. The shuttle vector p11559 was maintained using spectinomycin at concentrations of 70 µg/ml for *D. radiodurans* and 150 µg/ml for *E. coli*. Additionally, the shuttle expression vector pRADgro and its derivatives were maintained in the E. coli strain HB101, as previously described [35]. Standard molecular biology techniques were employed as described in the "Molecular Cloning: A Laboratory Manual" (4th Edition, Vol. II, Cold Spring Harbor Laboratory Press, New York, Green et al., 2012). Molecular biology-grade chemicals, enzymes, and salts were sourced from Sigma Chemicals Company (USA), Roche Biochemicals (Mannheim, Germany), New England Biolabs (USA), and Merck India Pvt. Ltd. (India). Radiolabeled nucleotides were provided by the Board of Radiation and Isotope Technology (BRIT), Department of Atomic Energy, India. For a comprehensive list of the bacterial strains and plasmids used in this study, please refer to Table S1.

### Cell survival studies

The experimental treatments for studying the survival of *D. radiodurans* cells involved exposure to various doses of UV and gamma radiation, as previously detailed [82], as well as treatment with MMC (10 µg/ml) according to the protocol described in a previous study [86]. Briefly, Bacterial cultures grown in TGY medium at 32°C were washed and suspended in sterile phosphate-buffered saline (PBS). These cultures were then exposed to varying doses of gamma radiation at a dose rate of 5.86 kGy per hour using a Gamma 5000 (^60^Cobalt) irradiator from the Board of Radiation and Isotope Technology, DAE, India. For UVC treatment, different dilutions of the cells were plated and subjected to 1500 J/m^2^ doses of UV radiation at 254 nm. The treated cells were subsequently plated on TGY agar plates, supplemented with appropriate antibiotics if necessary, and the colony-forming units were counted after a 48-hour incubation period at 32°C. For cells receiving both gamma radiation and MMC treatments, the protocol involves administering gamma radiation first, followed by MMC treatment. When combining UV with either gamma radiation or MMC, the sequence begins with gamma radiation or MMC treatment followed by UV exposure. For the spot assay, wild-type and NT gene mutants of *D. radiodurans* treated with appropriate DNA damaging agents alone or in combination of UV radiation cells diluted to appropriate serial dilution and 5 µl spotted on TGY plates supplemented with appropriate antibiotic wherever needed.

### Measurement of DNA repair kinetics using pulsed-field gel electrophoresis (PFGE)

Irradiated cultures were diluted in TGY to an OD600 of 0.2 and incubated at 30°C. At specified intervals, 5-ml samples were collected to prepare DNA plugs as described by Mattimore and Battista [129]. The DNA in the plugs was digested with 60 units of NotI restriction enzyme (Roche) for 16 hours at 37°C. Following digestion, the plugs underwent pulsed-field gel electrophoresis in 0.5x TBE using a CHEF-DR® III electrophoresis system (Bio-Rad) at 6 V/cm² for 20 hours at 14°C, with a linear pulse ramp of 50-90 seconds and a switching angle of 120°.

### Growth curve studies

*D. radiodurans* cells were cultured in TYG broth (containing 0.5% tryptone, 0.3% yeast extract, and 0.1% glucose) at 32°C under atmospheric conditions. The cultures were grown in a 24-well, energy-treated microplate (Nunc, Thermo Scientific™) for 16-20 hours using a BioTek Microplate Reader (Agilent Technologies, United States). For experiments requiring double-stranded cfDNA, a solution of 10 mg/ml deoxyribonucleic acid sodium salt from cfDNA (Sigma-D1501) was prepared and sonicated under sterile conditions. A final concentration of 0.5 mg/ml DNA was then added to either the minimal medium [130] or TYG medium as needed. The composition of the minimal medium for D. radiodurans was adopted from previously published sources (10831446).

---

**Table S1.**
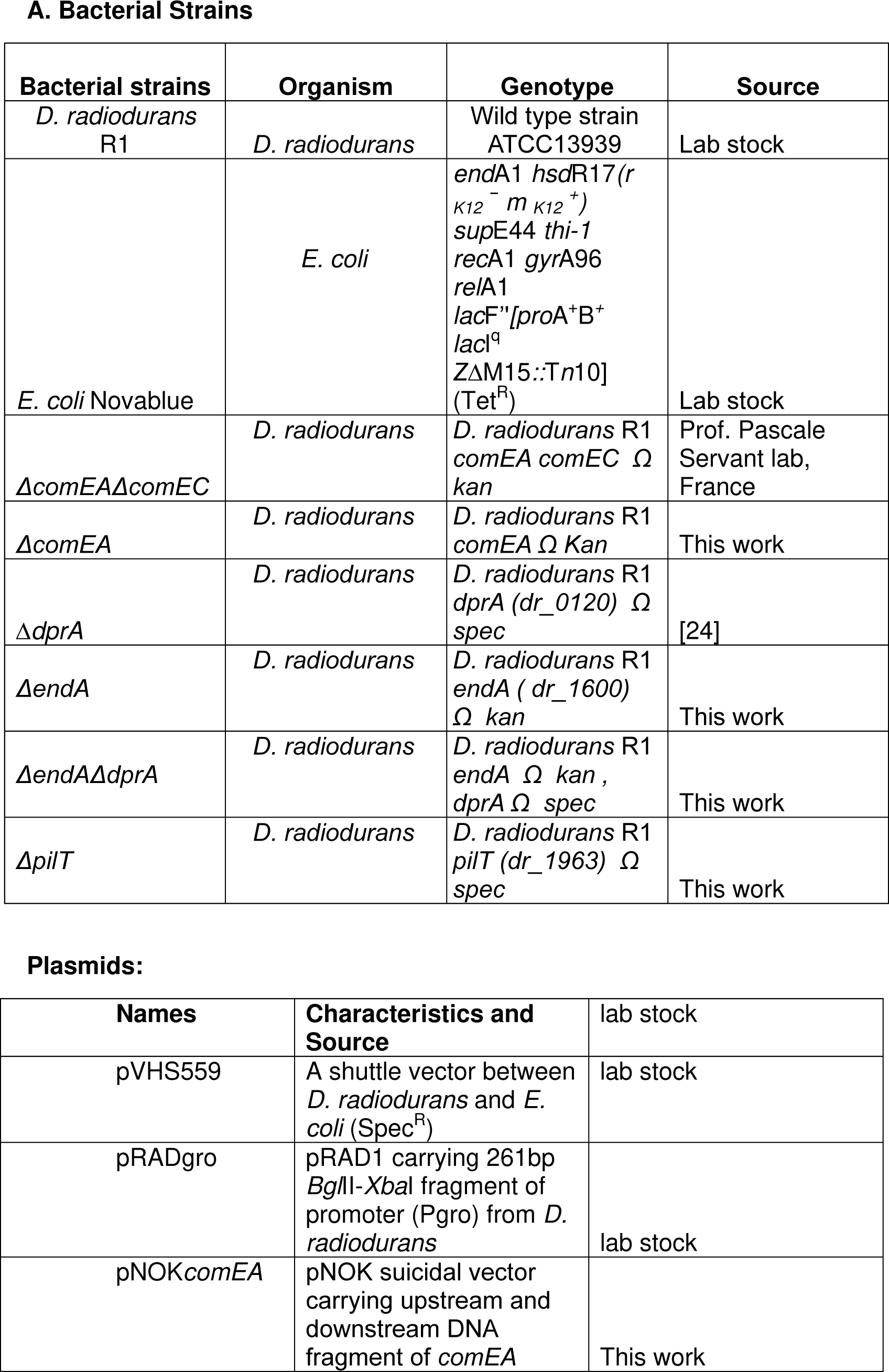
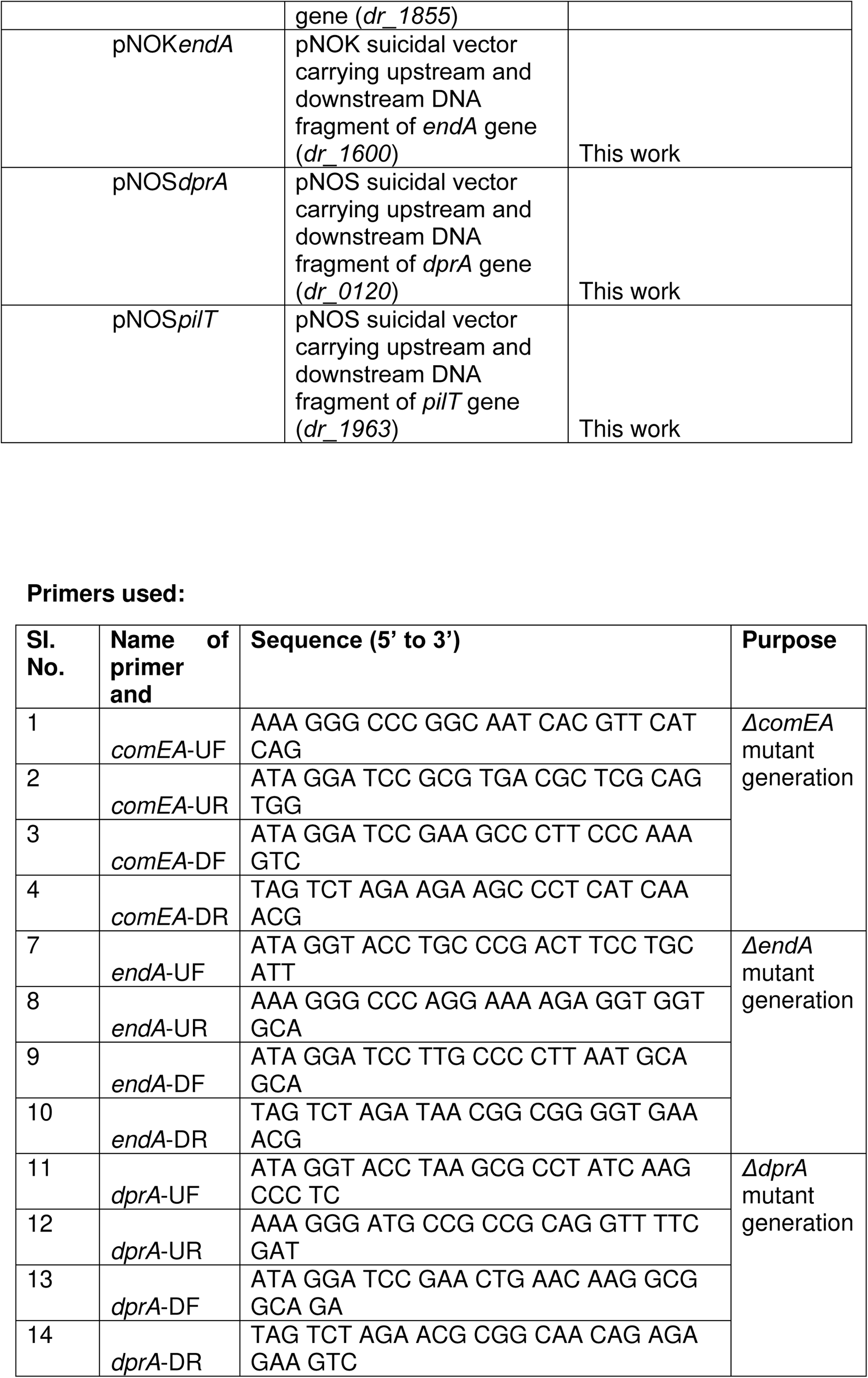
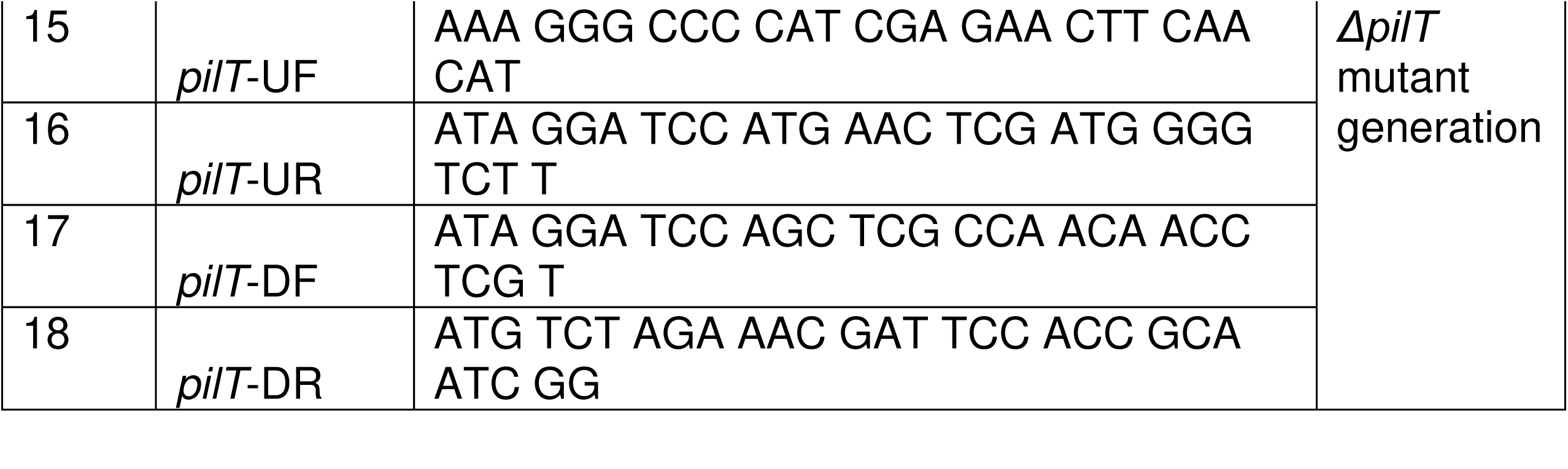
Bacterial strains, plasmids, and primers Used in this study.

